# Chronic high-sugar diet in adulthood protects *Caenorhabditis elegans* from 6-OHDA induced dopaminergic neurodegeneration

**DOI:** 10.1101/2023.05.29.542737

**Authors:** Katherine S. Morton, Jessica H. Hartman, Nathan Heffernan, Ian T. Ryde, Isabel W. Kenny-Ganzert, Lingfeng Meng, David R. Sherwood, Joel N. Meyer

**Affiliations:** Nicholas School of Environment, Duke University; Biochemistry and Molecular Biology, Medical University of South Carolina; Department of Biology, Duke University

**Author notes:** Declarations: The authors declare no competing interests.

**Keywords:** Glucose, Fructose, Neurodegeneration, Oxidative Stress, C. elegans

## Abstract

**Background:** Diets high in saturated fat and sugar, termed ‘western diets’, have been associated with several negative health outcomes, including increased risk for neurodegenerative disease. Parkinson’s Disease (PD) is the second most prevalent neurodegenerative disease and is characterized by the progressive death of dopaminergic neurons in the brain. We build upon previous work characterizing the impact of high sugar diets in *Caenorhabditis elegans* to mechanistically evaluate the relationship between high sugar diets and dopaminergic neurodegeneration.

**Results:** Adult high glucose and fructose diets, or exposure from day 1-5 of adulthood, led to increased lipid content and shorter lifespan and decreased reproduction. However, in contrast to previous reports, we found that adult chronic high-glucose and high-fructose diets did not induce dopaminergic neurodegeneration alone and were protective from 6-hydroxydopamine (6-OHDA) induced degeneration. Neither sugar altered baseline electron transport chain function, and both increased vulnerability to organism-wide ATP depletion when the electron transport chain was inhibited, arguing against energetic rescue as a basis for neuroprotection. The induction of oxidative stress by 6-OHDA is hypothesized to contribute to its pathology, and high sugar diets prevented this increase in the soma of the dopaminergic neurons. However, we did not find increased expression of antioxidant enzymes or glutathione levels. Instead, we found evidence suggesting downregulation of the dopamine reuptake transporter *dat-1* that could result in decreased 6-OHDA uptake.

**Conclusion:** Our work uncovers a neuroprotective role for high sugar diets, despite concomitant decreases in lifespan and reproduction. Our results support the broader finding that ATP depletion alone is insufficient to induce dopaminergic neurodegeneration, whereas increased neuronal oxidative stress may drive degeneration. Finally, our work highlights the importance of evaluating lifestyle by toxicant interactions.

## Background

In 2019 the average American consumed 50.0 g (190 calories) of refined cane or beet sugar and 29.3 g (111 calories) of high-fructose corn syrup per day in addition to other added caloric sweeteners and naturally occurring sugars (1). Despite a small 0.71% decrease in caloric sweetener consumption since 2016, this still often exceeds the World Health Organization recommendation of less than 10% of total caloric intake (2). Glucose and fructose are the most consumed sugars, as the majority of sugar intake in the United States comprises refined cane or beet sugar, high-fructose corn syrup, and foods naturally containing glucose and fructose (3). Referred to as high-glycemic-index diets for their propensity to raise blood glucose levels, high sugar diets have been linked to the increase in obesity with particularly strong evidence for the consumption of sugary beverages (4).

Obesity is defined as a body mass index (BMI) greater than 30 and is a non-monolithic disease caused by metabolic, genetic, socioeconomic, and environmental factors. It doubled in prevalence in more than 70 countries between 1980 and 2015 and is epidemiologically linked to the increased prevalence of type 2 diabetes, cardiovascular diseases, some neurodegenerative diseases, and surgical complications including infections (5, 6). In one such neurodegenerative disease, Parkinson’s disease (PD), it has been reported that patients show higher total sugar and added sugar consumption than healthy controls (7). Despite evidence of higher disease-concurrent intake in diagnosed individuals, it is less clear how sugar diets influence the onset of PD (8).

PD is a late-onset neurodegenerative disorder characterized by loss of function and death in the dopaminergic neurons of the substantia nigra region of the brain. PD impacts 1-3% of the global population over age 65. Oxidative stress and mitochondrial dysfunction have both been identified as potential causes or critical steps in the pathology of the disorder (9). Epidemiological studies consistently find relationships between blood glucose levels, insulin intolerance, and PD. Though the causal nature is unclear, increased blood glucose levels have been identified in drug-naïve patients and those showing cognitive decline (10), and elevated blood glucose is a predictor of cognitive decline (11). Increased added sugar intake has been associated with increased frequency of developing PD and greater symptom severity and medication requirements post diagnosis (12).These same studies and others have further found that decreased levels of insulin and increased insulin resistance were associated with cognitive decline in PD patients (10, 11, 13). It has not been clearly established if increased sugar intake and elevated blood glucose are causal or secondary to decreased insulin levels and sensitivity to insulin.

To expand understanding of the complex relationship between high sugar diets, obesity, and susceptibility to dopaminergic neurodegeneration, we turned to the nematode *Caenorhabditis elegans*. *C. elegans* has been widely used as a model in biomedical research in general and to explore the impacts of high-sugar diets in particular, because of its high genetic homology to humans, short life cycle, and conservation of key pathways including insulin signaling (14). *C. elegans* fed high-glucose diets generally demonstrate slower growth, decreased reproduction, shortened lifespan, neuronal and mitochondrial dysfunction, decreased anoxia survival, and increased oxidative stress (15–19). High-fructose diets, though less explored, have been shown to decrease lifespan and health span, induce mitochondrial swelling, and decrease anoxia survival of worms (19, 20). *C. elegans* has also been employed for studies of PD. Possessing 8 dopaminergic neurons and high genetic tractability, several transgenics have been generated to assist with visualization of dopamine neuron morphology (21, 22). High glucose exposure studies in *C. elegans* showed increased susceptibility to organophosphate pesticide-induced neurodegeneration in dopaminergic, GABAergic, and cholinergic neurons (17, 18, 23). These studies, however, were mostly performed with acute, developmental exposures to glucose.

Here, we present evidence from worms fed chronic, not acute, 100 mM D-glucose or fructose from day 1-day 5 of adulthood on the mechanistic relationships between high-sugar diets and dopaminergic neurodegeneration. With this strategy, we have avoided the potential for confounding effects of bioenergetic remodeling resulting from developmental mitochondrial stress (24–28). Doses were selected to closely match the large body of literature in *C. elegans* studying high sugar diets, in which 100 mM consistently produces clear effects but is non-lethal (Table 1). To improve upon the common use of decreased fluorescence of the cell bodies within the cephalic (CEP) neurons in the head of the worm as a proxy for degeneration, we employ a neurodegeneration scoring methodology with improved ability to detect subtle changes to the neuronal processes. We not only assess whether high sugar diets induce dopaminergic neurodegeneration, but whether they enhance susceptibility to the canonical dopaminergic neurotoxicant 6-hydroxydopamine (6-OHDA). Upon entering the cell, 6-OHDA increases oxidative stress partly through auto-oxidation, and partly through inhibition of mitochondrial electron transport chain Complexes I and IV, resulting in decreased ATP levels, akin to two other PD toxicant model toxicants, rotenone and MPP+ (29). Utilizing this method, we report that chronic, adult high glucose or high fructose diets resulted in neuroprotection from 6-OHDA exposure. The impact of 6- OHDA on redox state, not its effect on ATP levels, was abrogated by high sugar, suggesting that redox alterations, not energetic alterations, underlie the dopaminergic neurotoxicity of 6-OHDA in *C. elegans*. In absence of alterations to glutathione levels, redox tone, and antioxidant enzyme expression, we suggest altered dopamine neurotransmission leads to decreased 6-OHDA uptake and prevents toxicity.

**Table 1:**
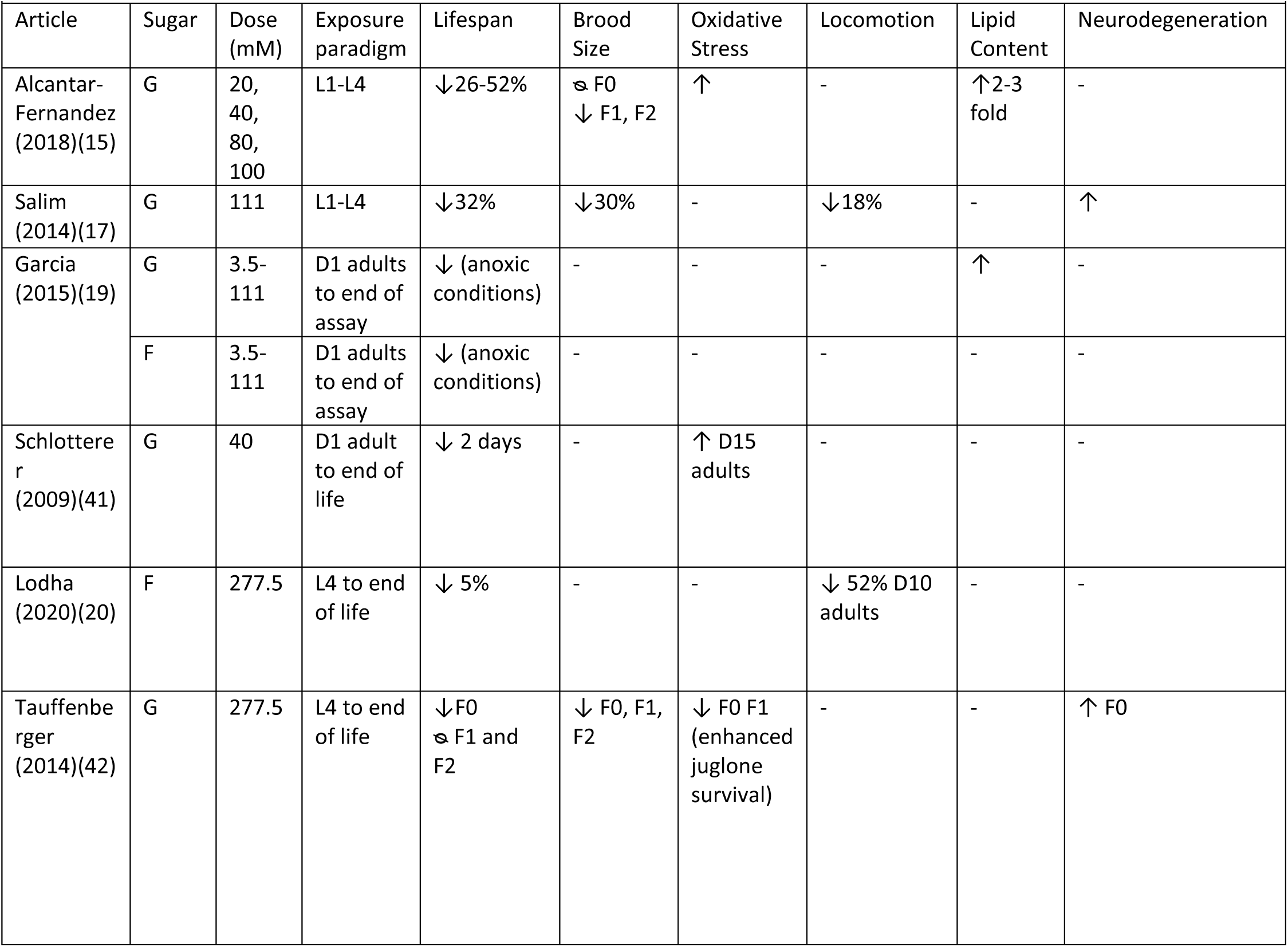
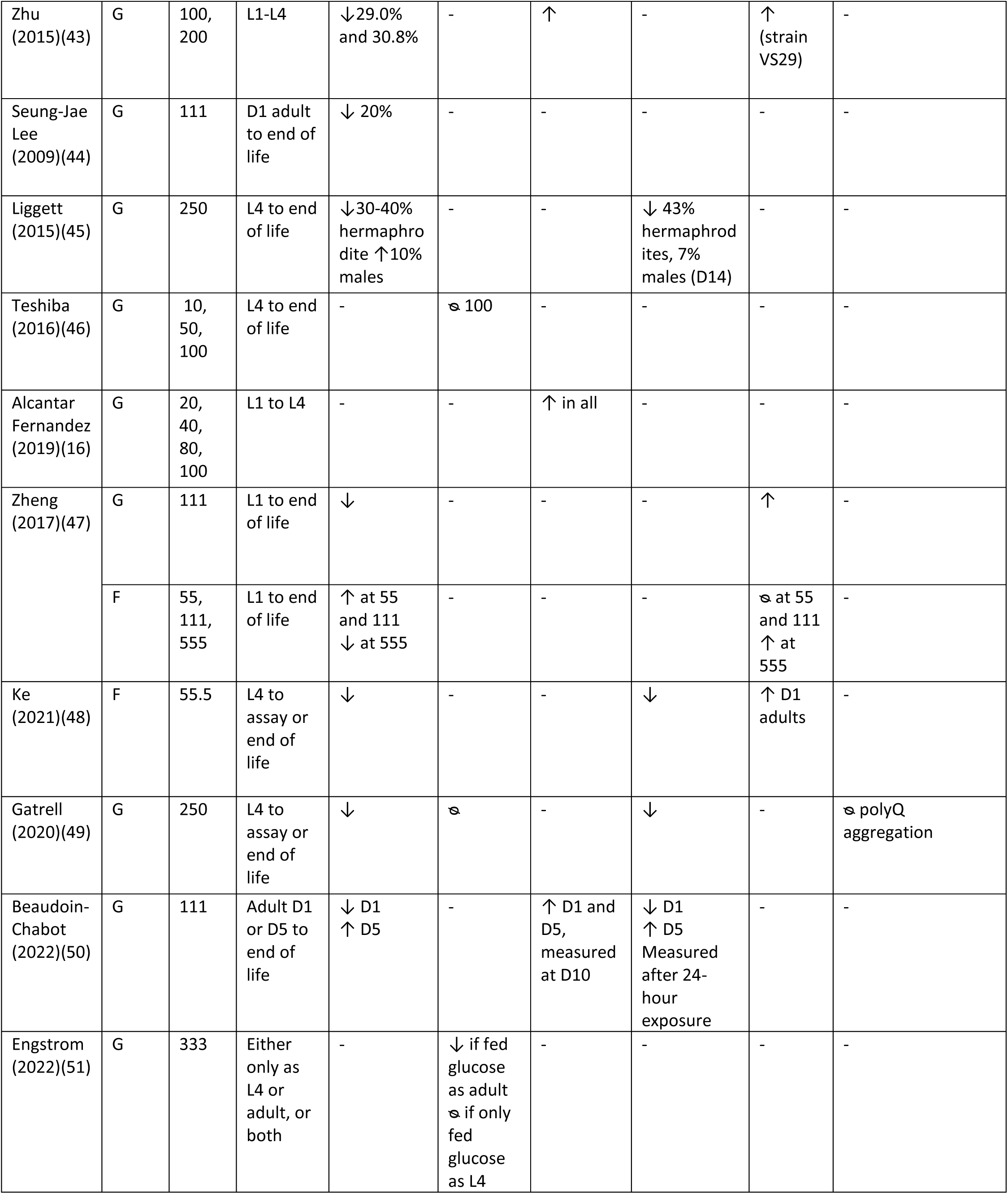
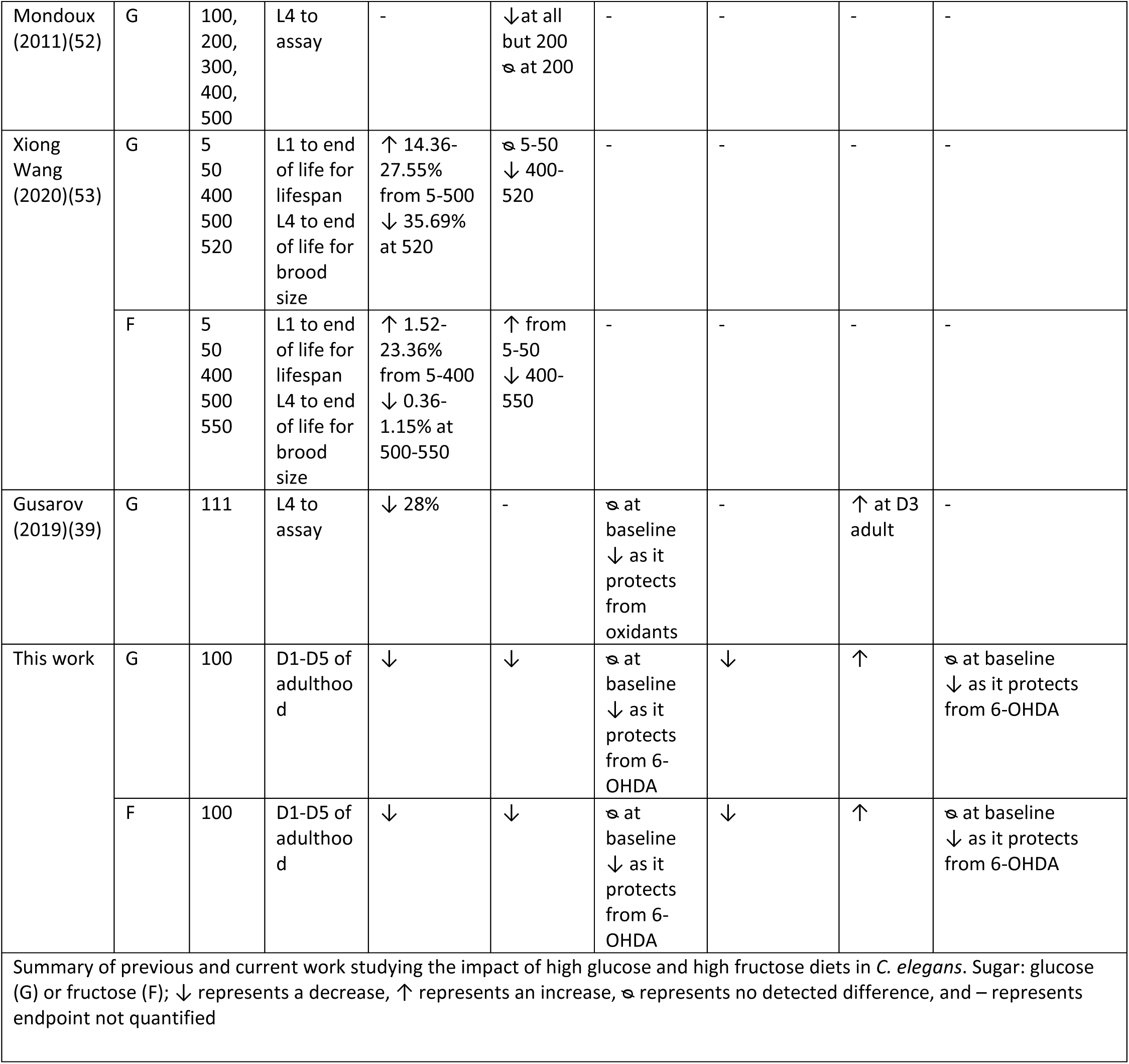
Summary of previous research on high glucose and high fructose diets in *C. elegans*.

## Results

### Adult high sugar diets decrease lifespan and fecundity while increasing adiposity

*C. elegans* has been used extensively to evaluate the impacts of dietary paradigms on lifespan and reproduction. In the case of high sugar diets, previous work in *C. elegans* has focused largely on exposures beginning in early development. To discern how adult glucose and fructose exposures impact key biological functions, and permit comparison to previously published developmental exposure studies, we first evaluated the impact of our adult exposure paradigm on adiposity, lifespan, and fecundity. In concurrence with the effects observed with developmental sugar exposures, worms transferred as young adults to nematode growth media (NGM) plates supplemented with 100 mM glucose or 100 mM fructose show increased lipid accumulation represented by an 85.6% and 46.2% increase in fluorescence intensity of an mCherry::mdt-28 fusion protein localized primarily to lipid droplets (Fig 1A). Increased lipid stores were particularly concentrated throughout the intestine, around the vulva, and to a lesser extent in the head. The effect was more intense in glucose fed worms than fructose fed worms (Fig 1B). Similarly, and again in agreement with observations from developmental exposures, worms fed glucose and fructose had a modest decrease in their average brood size from 297.7±7.9 eggs per worm to 247.9±7.9 and 272.7±4.8 respectively (Fig 1C). Beyond total brood size, the time course of egg laying was altered such that both high sugar diets caused egg laying to be distributed more evenly over days 1-3 of adulthood as opposed to most being laid the first 2 days with a sharp decrease on the third (Fig 1D), which is the typical pattern on control plates. Finally, the sugar exposure paradigm shows no significant lethality during exposure, but both sugars led to a significantly decreased median lifespan post exposure (SFig 1, Fig 1E), with a greater decrease on fructose (6 days shorter than control diet) than glucose (2 days shorter).

**Figure 1.**
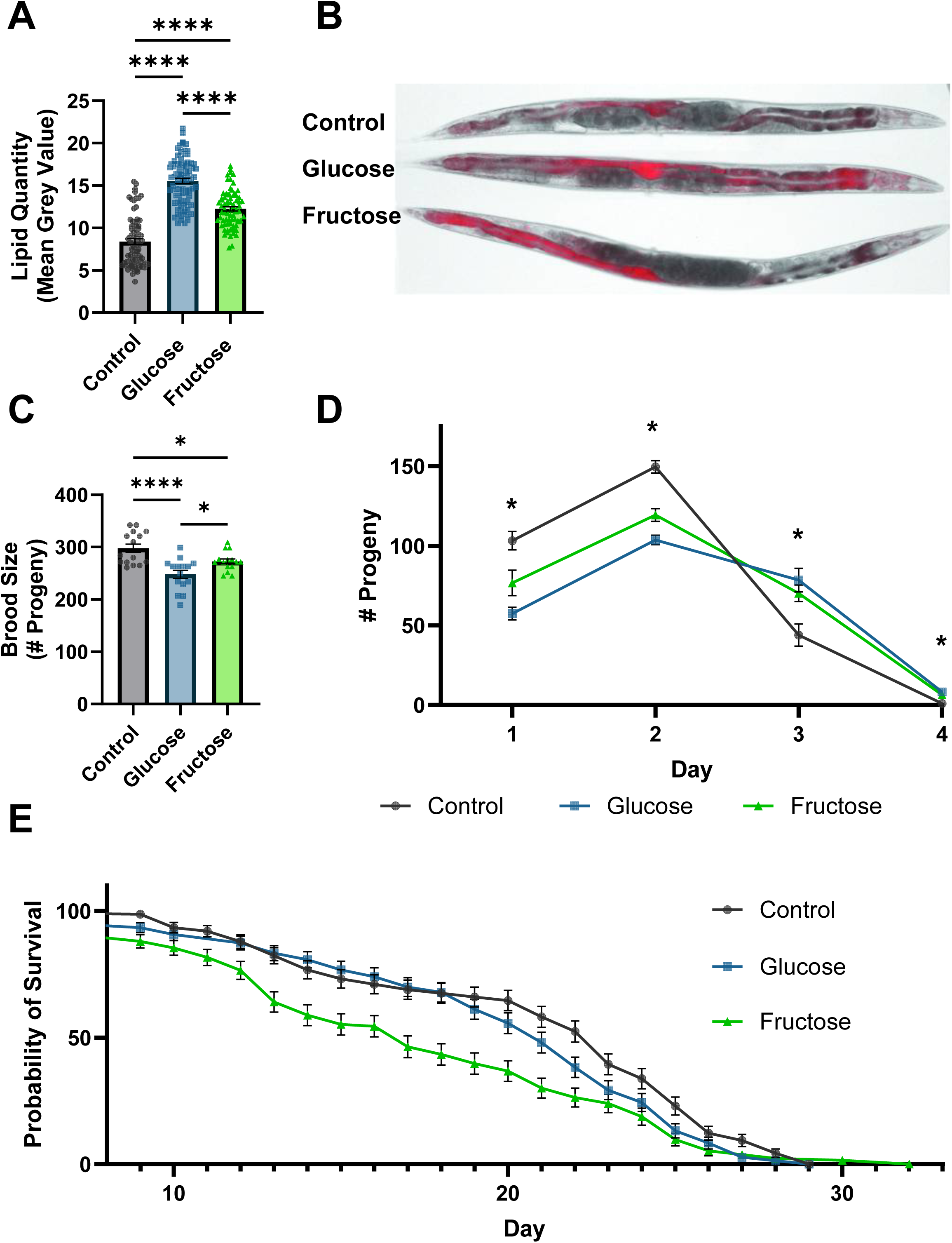
High-sugar diets increase lipid content, decrease reproduction, and shorten lifespan. **A** Fat quantification and **B** representative images of 8-day old LIU2 worms reared on control (n=77),100 mM glucose (n=83), or 100 mM fructose (n=74) supplemented NGM plates with OP50 as a food source. **C** Total number of progeny (n=15 per treatment) **D** progeny laid per day (n=45, 15 per treatment, p-interaction<0.0001) of individual worms plated on 6 cm control or sugar supplemented plates from late L4 to post-reproductive age. **E** Lifespan analysis of worms treated from days 1-5 of adulthood on control (n=141, median survival 23 days), glucose (n=146, median survival 21 days, p=0.0271), or fructose (n=137, median survival 17 days, p=0.0001) supplemented NGM plates then transferred to K-agar OP50 plates until death. Only worms that were alive on day 5 of adulthood were utilized. For **A-D**, Three biological replicates were performed for each experiment. Shapiro Wilks normality tests were used to confirm distribution normality of the data. One-way ANOVA followed by Tukey’s Post-Hoc was used for A and C to determine p-value. For D, a two-way ANOVA with Tukey’s post-hoc was used. For **E**, a Kaplan-Meier survival analysis was performed in conjunction with the Log-rank test. *p<0.0332, **p<0.0021, ***p<0.0002, ****p<0.0001

### High sugar diets protect from 6-OHDA induced dopaminergic neurodegeneration

To define the role of high sugar diets in age-related and toxicant-induced neurodegeneration, we compared dendritic degeneration in worms exposed to high sugar diets throughout reproductive adulthood and subsequently exposed to either 25 mM or 50 mM 6-hydroxydopamine (6-OHDA). 6- OHDA is a well-validated dopaminergic neurotoxicant transported into the dopaminergic neurons via the DAT-1 transporter. The CEP neurons in *C. elegans* are easily visualized within the head of the worm and have a well characterized damage phenotype including dendritic blebbing and breaking (21, 22, 30–32). Using a qualitative scale in which increasing score represents increasing damage, we found that high sugar diets did not increase age related neurodegeneration. In response to 25 mM 6-OHDA exposure high glucose was protective, and both sugars were protective at the 50 mM dose with fewer instances of broken and fully deteriorated sections of dendrite (Fig 2A, 2B, SFig 2).

**Figure 2.**
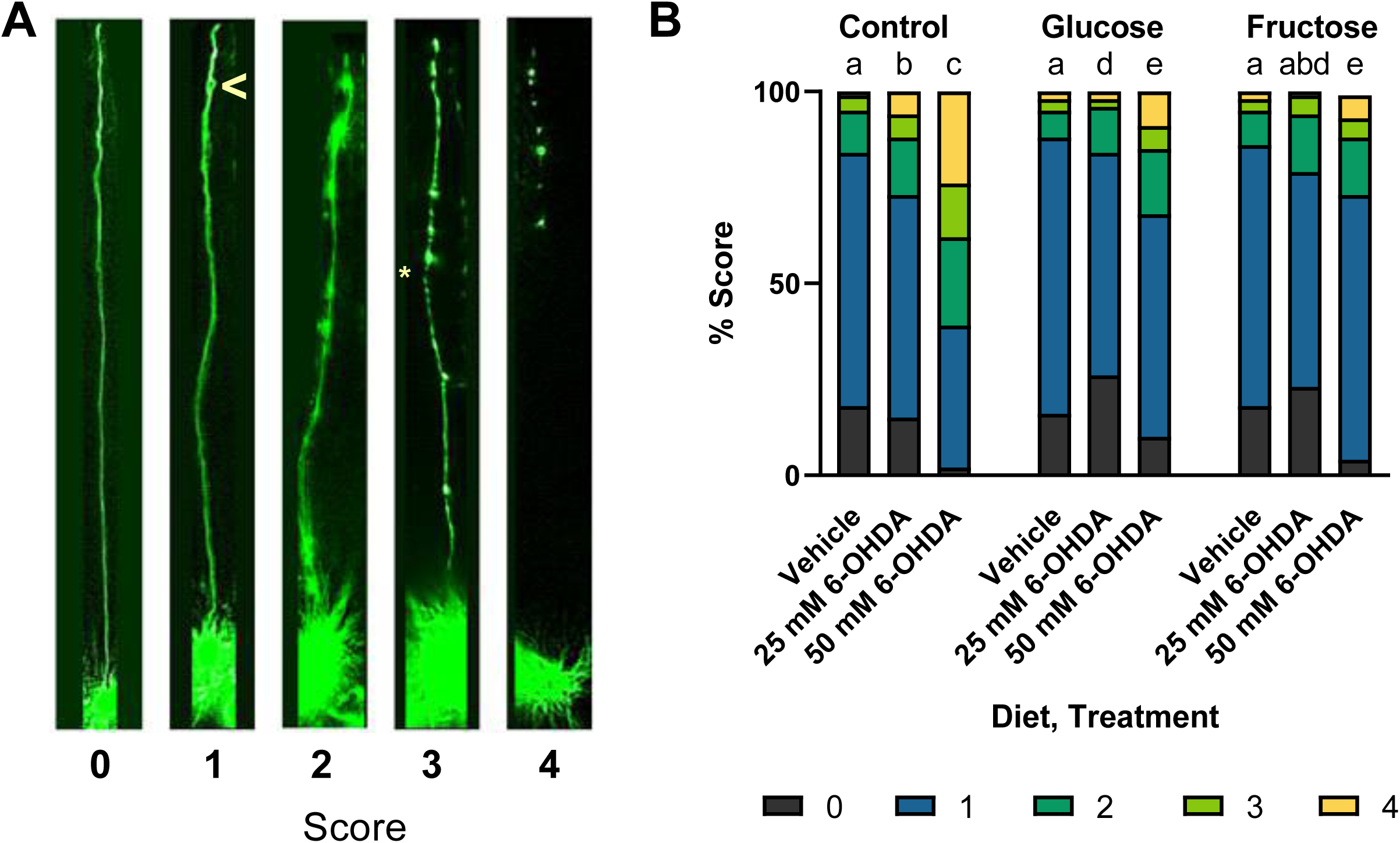
High-sugar diets do not induce neurodegeneration and protect from the neurotoxicant 6- hydroxydopamine (6-OHDA). **A** Representative images for each of the 5 scores used to assess dopaminergic neurodegeneration. The “>” symbol in the score of a 1 denotes a bleb, and the “*” in the score of a 3 denotes a break. **B** A comparison of neurodegeneration in control, glucose-fed, and fructose-fed worms treated with a vehicle control of 5 mM Ascorbic Acid or 25 mM or 50mM of 6- OHDA. Pairwise chi-squared analysis was run with a Bonferroni corrected p-value of < 0.003571 to account for 14 pairwise comparisons. Statistical difference is represented by letters a-e such that bars possessing the same letter are not statistically different, and bars possessing none of the same letters are statistically different. Data from 6 biological replicates is represented for total n=2,676, n per group =172-512.

### Neuroprotection by high sugar diets is not explained by alterations in mitochondrial amount or morphology

Next, we worked to understand the mechanism for the observed glucose and fructose neuroprotection. As the 6-OHDA exposure paradigm is acute (one hour), we reasoned that the mechanism of protection resulting from the chronic high sugar diet must be present when the exposure begins. Both high glucose and high fructose diets have been previously associated with increased mitochondrial swelling and fragmentation (20, 33). Increases or decreases in mitochondrial fission and fusion dynamics are vital to cellular response to dietary and toxicant exposures (34, 35). Mitochondrial fission is required for the increase in reactive oxygen species and induction of cell death by high glucose diets in some cell types (36). Therefore, we evaluated mitochondrial morphology and number in the CEP neuron dendrites and mitochondrial area in muscle cells to determine if clear differences in mitochondrial dynamics were present prior to 6-OHDA exposure. No differences were apparent in the muscle cell mitochondrial area (SFig 3) or in neuronal cell mitochondrial number (Fig 3A); however, high glucose diets altered mitochondrial morphology within the CEP neurons (Fig 3B-3D). In CEP neurons, high glucose diets resulted in small but significant elongation of the mitochondria from 1.832±0.04 micrometers to 2.005±0.05 micrometers in length (Fig 3A-B). Similarly, the maximum length of mitochondria increased from 3.98±0.11 to 4.50±0.16 (Fig 3C). This resulted in an increase of total mitochondrial length per dendrite from 16.45±0.32 micrometers to 18.64±0.45 micrometers (Fig 3D).

**Figure 3.**
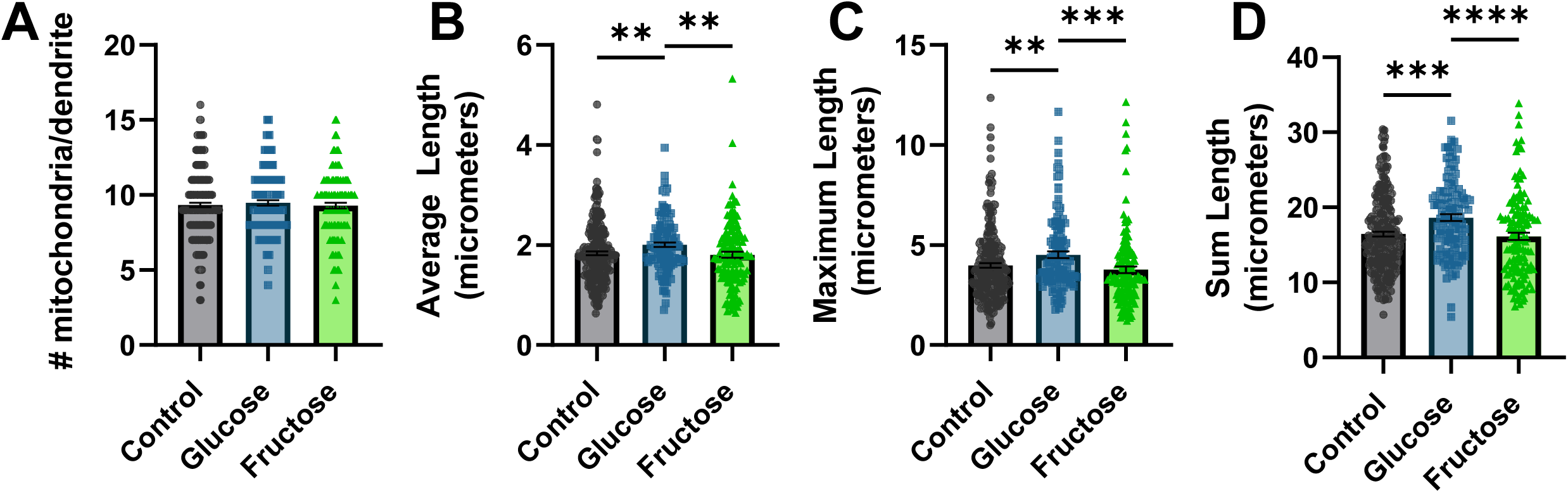
High-glucose diet induces mild neuronal mitochondrial elongation. **A** The number of mitochondria per dendrite, **B** average length of mitochondria per dendrite, **C** length of the longest mitochondria per dendrite, and **D** sum of the lengths of all mitochondria within a dendrite in worms reared on control (n=249), 100 mM glucose (n=127), or 100 mM fructose (n=134) supplemented NGM plates. **A-D** Three biological replicates were performed. Shapiro-Wilks Normality tests determined all data sets were non-normally distributed. Kruskal-Wallis test followed by Dunn’s multiple comparisons testwas used to establish p-values. *p<0.0332, **p<0.0021, ***p<0.0002, ****p<0.0001

However, high-fructose diets did not result in any significant alterations to mitochondrial number or morphology within the dendrites, indicating that even if glucose-mediated increased mitochondrial length contributed to protection from 6-OHDA, fructose’s protective effect could not be explained by this mechanism. Continuing to assess mitochondrial mechanisms that could confer protection, we moved to evaluate cellular and organismal bioenergetics.

### High sugar diets do not rescue ATP depletion caused by electron transport chain inhibition

It has been theorized that ATP depletion may incite a negative feedback loop resulting in and enhancing neurodegeneration (37, 38). Therefore, we next assessed whether the high sugar diets protected from dopaminergic neurodegeneration by improving energetics at baseline, or upon challenge. On an organismal level, we assessed mitochondrial bioenergetic function by whole worm respirometry. We found a small increase in basal oxygen consumption rate (SFig 4); however, this is accounted for by larger worm size. After accounting for worm size, we found no alterations to electron transport chain function or non-mitochondrial oxygen consumption (Fig 4A). We also assessed whole-worm ATP levels by luminescent assay and observed no baseline differences (Fig 4B). To assess energetic status upon challenge, we exposed worms to the complex I inhibitor rotenone. This acute (1-hour) challenge decreased whole-body ATP levels of sugar-fed worms 40% more than controls (Fig 4B). To determine if energetic responses in the CEP neurons follow the same trend as the whole-organism responses, and assess whether that susceptibility would manifest in the context of the 6-OHDA challenge that we used for neurodegeneration, we exposed worms expressing the PercevalHR ATP:ADP ratio reporter in dopaminergic neurons to 50 mM 6-OHDA and vehicle controls of ascorbic acid. Surprisingly, ATP:ADP ratio within the CEP neuron soma was not different across diets before or after 6-OHDA exposure (Fig 4C). Notably, ascorbic acid, the vehicle for 6-OHDA, induced significant ATP depletion. As ascorbic acid does not induce neurodegeneration, and both sugars protected from neurodegeneration without protecting from ATP depletion, our results are inconsistent with ATP depletion causing degeneration of the CEP neurons.

**Figure 4.**
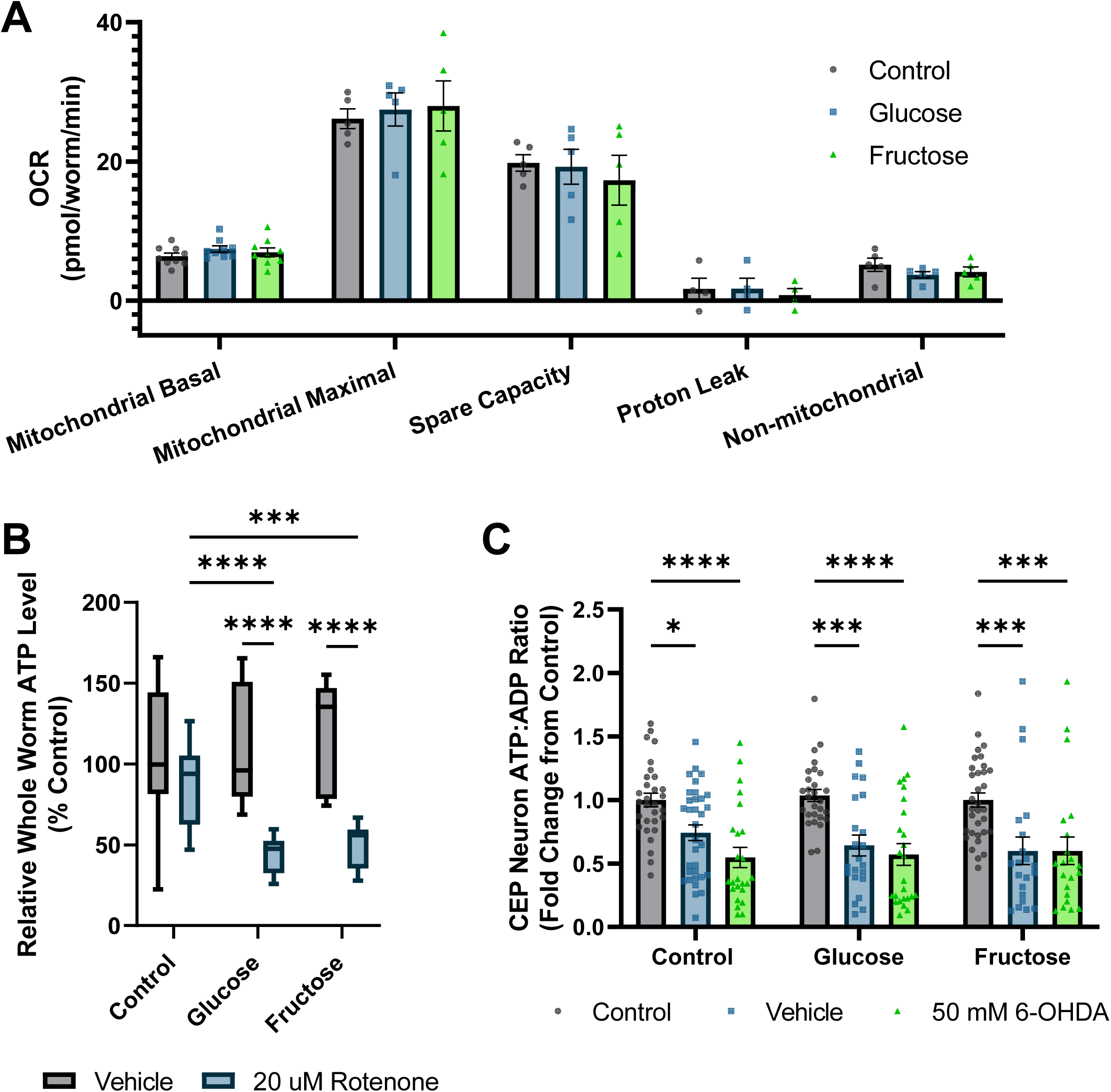
High sugar diets do not alter baseline bioenergetics or protect from ATP depletion from mitochondrial inhibitors. **A** Whole worm respirometry was performed on D8 worms after high-sugar diet exposure to quantify mitochondrial respiratory function. Oxygen consumption rate was normalized to worm number and volume to account for differences in body size. **B** Whole worm ATP levels were quantified after dietary exposure to either control, high-glucose, or high-fructose conditions. Control, glucose, and fructose exposed worms were also subjected to a 1-hour 20 uM rotenone challenge to assess organismal response to electron transport chain inhibition. **C** Worms expression dat- 1::PercevalHR were exposed to control, high-glucose, or high-fructose conditions, and assessed on day 8 under after exposure to ascorbic acid or 50 mM 6-OHDA. *p<0.0332, **p<0.0021, ***p<0.0002, ****p<0.0001

### High sugar diets minimally alter organismal antioxidant gene expression and do not change glutathione concentrations

The third mechanism we tested for sugar-mediated dopaminergic neuroprotection was upregulation of antioxidant defenses. Acute high sugar diets have been demonstrated to increase oxidative stress; however, more chronic exposure in young adults increased expression of the proteins glucose-6- phosphate 1-dehydrogenase and glutathione disulfide reductase, which should allow for accelerated reduction of glutathione and confer protection from oxidant exposures (39). We hypothesized that chronic high-sugar diets might cause similar compensatory and protective upregulation of antioxidant systems, which could combat redox stress induced by 6-OHDA exposure. First, we assessed if our chronic high sugar diets altered organismal redox state using a whole-animal reduction oxidation sensitive GFP (roGFP) construct that reports on the ratio of oxidized to reduced glutathione. Due to the increase in autofluorescence at 405nm driven primarily by gut autofluorescence (Supp Fig 5A-B), we restricted our analysis to the head region from the tip of the head of the worm to the end of the terminal pharyngeal bulb. High sugar diets induced no differences in the redox tone of the glutathione pool on an organismal level (Fig 5A). To ensure that the lack of alteration in glutathione redox state was not due to differences in total glutathione pool sizes, we quantified total glutathione levels and found no statistically significant difference (Fig 5B). To determine if this result, which was contrary to findings from acute exposures, was due to altered antioxidant gene expression, we evaluated the mRNA expression levels of multiple antioxidant enzymes. We observed slight (10-20%) decreases in expression of the glutathione reductase encoding gene *gsr-1* in glucose fed worms, and of a cytosolic CuZnSOD encoding gene, *sod-5,* in fructose fed worms (Fig 5C, SFig 6). Decreased expression of antioxidant genes may lead to susceptibility to oxidant exposures, leading us to examine redox state specifically within the mitochondria of the CEP neurons that were targeted by 6-OHDA in our neurodegeneration studies. We utilized a CEP neuron-specific mitochondrial-targeted roGFP and, in accordance with the organismal result, no difference was observed as a result of the high sugar diets alone. However, sugar-fed worms had a significantly smaller increase in oxidation state after 6-OHDA exposure (Fig 5D). This result is consistent with protection from neurodegeneration and of oxidative stress as a driver of neurodegeneration. However, it could be explained either by a cell-specific increase in antioxidant defenses, not detectable by our whole-organism measurements, or by a decrease in 6-OHDA uptake by the dopaminergic neurons. We next tested the latter possibility.

**Figure 5.**
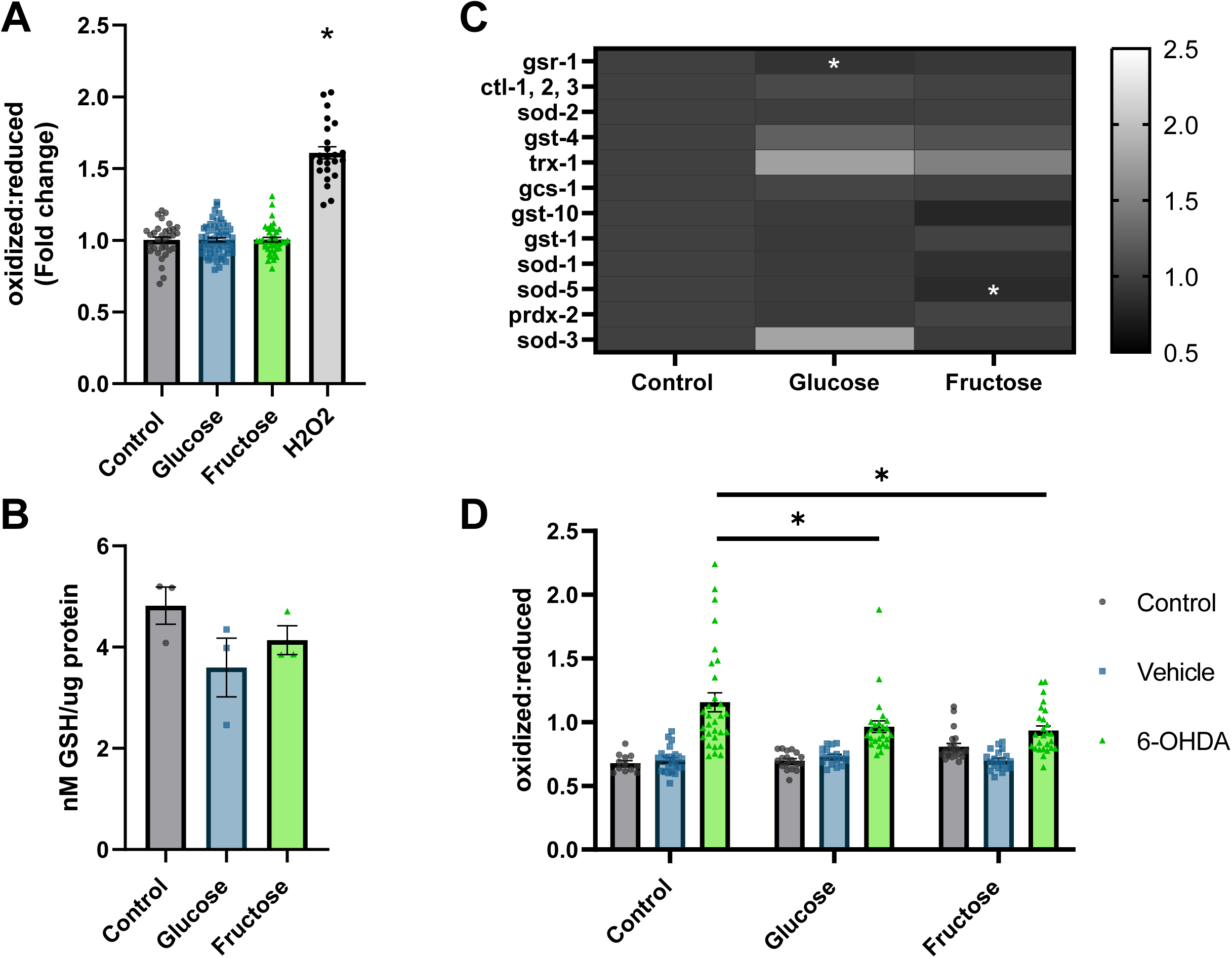
High-sugar diets protect from 6-OHDA induced oxidative stress minimal alteration to antioxidant systems. **A** The redox tone of the glutathione pool was quantified to assess organismal oxidative stress. Worms expressing reduction:oxidation sensitive GFP were reared from day 1-5 of adulthood on NGM plates or NGM supplemented with 100 mM glucose or fructose. Control worms were exposed to 3% H2O2 as a positive control for increased oxidation. Three biological replicates were assessed with n: Control=33 Glucose=58 Fructose=40 H_2_O_2_=24 **B** Organismal total glutathione levels were assessed control, glucose, and fructose fed worms. Three biological replicates were utilized with 2- 3 technical replicates averaged to produce n=1 per biological replicate. **C** Alterations to mRNA levels of multiple families of antioxidants were assessed for alterations by qPCR. Three biological replicates were performed with n=3 per replicate, total n=9 per treatment. **D** To examine redox response within the CEP neurons to 6-OHDA, worms expressing *dat-1*::mls roGFP were used and exposed to control, vehicle (5 mM Ascorbic Acid) or 50 mM 5-OHDA on day 5 of adulthood. Three biological replicates are represented with total n=195 **A-B** Normality was confirmed by a Shapiro-Wilks normality test; One-way ANOVA followed by Tukey’s Post-hoc was used to determine p-values. **C** Relative gene expression was determined by the ΔΔCt method, and each gene was analyzed by one-way ANOVA with Tukey’s Post- hoc. **D** One-way ANOVA followed by Tukey’s Post-hoc was used to determine p-values. For all panels *p<0.0332, **p<0.0021, ***p<0.0002, ****p<0.0001

### High sugar diets modulate the dopamine transport system to decrease dopamine reuptake

6-OHDA is actively transported into the dopaminergic neurons by the DAT-1 transporter. Each worm has only 4 CEP neurons, which makes direct quantification of the uptake of 6-OHDA impractical. Therefore, we instead measured proxies for the quantity and activity of the DAT-1 transporter. We assessed mRNA expression of *dat-1*, the re-uptake transporter responsible for 6-OHDA transport into the CEP neurons; *cat-1*, a vesicular monoamine transporter critical to dopamine packaging and release (40); *cat-2*, which encodes tyrosine hydroxylase, the protein that catalyzes the rate limiting step in dopamine synthesis; and the dopamine receptor *dop-3* (Fig 6A). With no alterations in expression levels observed via rtPCR, we also examined DAT-1 expression via a *dat-1* promoter driven GFP strain. We detected a 28.16±3.30% and 26.03±3.15% decrease in *dat-1* promoter-driven fluorescence in glucose and fructose exposed worms, respectively (Fig 6B). To further evaluate the functional status of dopaminergic neurotransmission, we utilized a swimming induced paralysis (SWIP) assay. Release of dopamine into the neuro-muscular junction dictates the ability of the muscle cells in worms to contract and relax.

**Figure 6.**
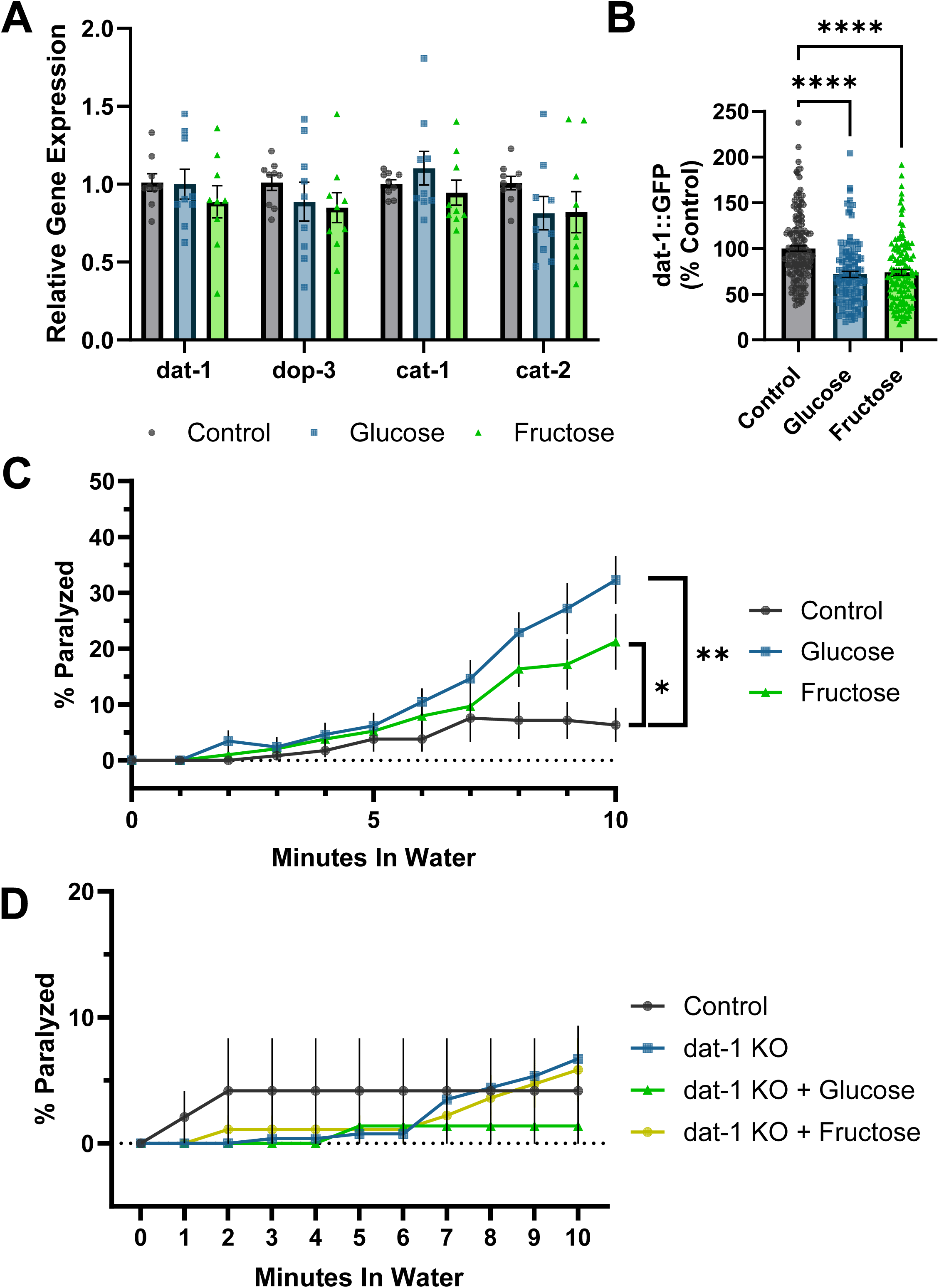
Dopamine transmission is altered in sugar-fed animals. **A** Relative expression of dopamine synthesis, reuptake, and transporter genes was assessed via qPCR in worms reared on control NGM or high-glucose or high-fructose supplemented plates. A one-way ANOVA with Tukey’s post hoc was used to determine p-values. **B** Quantification of pdat-1::GFP fluorescence intensity was assessed as a broader measure of dat-1 transcription. Results represent three biological replicates, n=157, 112, 129 respectively. These data are non-normally distributed (Shapiro-Wilk’s Normality p<0.05) and were evaluated by a Kruskal-Wallis test, ****p<0.0001. **C** The tendency of sugar fed worms to undergo swimming induced paralysis (SWIP) was determined. A two-way ANOVA with Dunn’s post hoc was used to assess significance. Four biological replicates are represented, n=12 for each diet. *p<0.0380, **p<0.0002 **D** The tendency of *dat-1* KO worms to undergo SWIP compared to BY200 controls under control and high-sugar diet conditions. Three biological replicates are represented, n=6 for each treatment.

When too much dopamine enters the junction, or not enough is cleared via re-uptake, the worms are temporarily paralyzed. After 10 minutes of swimming, glucose and fructose fed worms were five and two times more likely to SWIP (Fig 6C). Increased SWIP activity may indicate decreased dopamine reuptake by the CEP neurons, which would protect against 6-OHDA uptake. To confirm the role of *dat-1* in this phenotype, we performed the SWIP assay with worms possessing a 1836 base pair knock out (KO) in *dat-1*. Unlike their response in early life, day 8 *dat-1* KO worms do not SWIP more than controls, implying an adaptive response throughout life (SFig 7, Fig 6D). However, *dat-1* KO worms fed glucose and fructose are also not susceptible to SWIP, supporting our hypothesis that DAT-1 downregulation or internalization as a result of high sugar diets is the source of elevated susceptibility of sugar fed worms to SWIP (Fig 6D). Together, these data support the overarching hypothesis that chronic sugar mediated *dat-1* downregulation decreases 6-OHDA induced dopaminergic neurodegeneration.

## Discussion

Sugars such as glucose and fructose are essential for animal life, but diets containing excessive sugar can increase neurodegeneration in mammalian models and *C. elegans* (18, 23, 49, 54). We expand on previous investigations of high sugar diets in *C. elegans* to investigate mechanistic links between high sugar in adulthood, mitochondrial disfunction, and dopaminergic neurodegeneration (Table 1). Although our exposure paradigm begins in early adulthood, like previous work, it still led to decreased lifespan, increased lipid accumulation, and decreased reproduction. Notably, the rapid onset of reproductive changes is cohesive with previous reports demonstrating adult exposure leads to decreased progeny, while beginning exposure at late-L4 likely drives the slowed time course of reproduction by altering germline proliferation, meiotic entry, or sex differentiation(51). Despite these similarities, our results were inconsistent with previous findings in which high glucose induced degeneration of dopaminergic neurons, decreased dopamine levels, and exacerbated monocrotophos-induced neurotoxicity (18, 55). In this study we did not find a change in neurodegeneration after chronic, adult high-glucose and high-fructose diets. Rather, we found that these diets protected from 6-OHDA-induced dopaminergic neurodegeneration. After assessing a number of potential mechanisms of protection, we propose that the protective effect is mediated by decreased 6-OHDA uptake via the DAT-1 transporter.

Contrary to previous work showing electron transport chain impairment and severe mitochondrial dysfunction (16, 20, 54), we only identified a slight elongation of the mitochondria in dopaminergic neurons of glucose fed worms, and no change in those fed fructose. No difference in mitochondrial function was detected with whole worm respirometry, total ATP levels, or ATP:ADP ratio within the CEP neurons. The only apparent indication of bioenergetic dysfunction induced by the high sugar paradigm was in response to challenge by the Complex I inhibitor rotenone, which caused nearly 40% greater ATP depletion in sugar-fed worms. 80 mM and 100 mM glucose dose-dependently decreased the activity of Complex I in a previous *C. elegans* study, without alteration to ATP, ADP, or AMP concentrations (16). It is plausible that lower Complex I activity decreased the dose of rotenone required to completely inhibit Complex I function, or that glycolysis is already enhanced by the high sugar diets, preventing further transition to glycolytic metabolism. We previously demonstrated the upregulation of glycolysis in rotenone-treated worms (56), though not in the context of high-sugar diets. Because different cell types rely on different bioenergetic pathways, we next tested whether the increased susceptibility to acute electron transport chain inhibition we observed on the organismal level would also be observed within the CEP neurons. Remarkably, sugar-fed worms showed no discernable difference in ATP:ADP ratio after 6-OHDA exposure, which is inconsistent with energetic deficit causing neurodegeneration, since the same sugar exposures protected against 6-OHDA-induced neurodegeneration. Furthermore, as the vehicle in our neurodegeneration experiments also decreases ATP:ADP ratio but did not cause neurodegeneration, it is improbable ATP depletion is the mechanistic step responsible for 6-OHDA induced dopaminergic neurodegeneration. Earlier work characterizing rotenone similarly noted that redox stress, not ATP depletion, is critical for its induction of neurodegeneration (57, 58).

Previous studies with acute sugar exposure models have detected increased antioxidant enzyme expression or increased total glutathione levels (15, 39), which would protect from the oxidative stress induced by 6-OHDA. However, we found no large differences in total glutathione, redox tone of the glutathione pool, or expression of antioxidant enzymes in the glutathione-related, superoxide dismutase, catalase, peroxiredoxin, or thioredoxin families. It is possible that differences in mRNA and/or protein levels specifically within the CEP neurons existed but were not detected in our whole-organism gene expression analysis. However, many previous studies that detected upregulated antioxidant defenses after acute exposures also employed whole-organism measures, making this explanation less likely. Perhaps more likely, our chronic exposure paradigm may result in adaptations across the lifetime of the worm in glucose uptake, transport, and utilization, culminating in a loss of the acute-phase oxidative stress response, explaining our lack of effects. Thus, despite finding a decreased redox response to 6-OHDA in the CEP neurons of sugar-fed worms, this resilience to oxidative challenge is unlikely a result of enhanced antioxidant defenses. Having failed to find compelling evidence for redox changes or bioenergetic inhibition as the mechanism for neuroprotection, we next considered the possibility of altered 6-OHDA uptake in high-sugar fed worms.

The DAT-1 transporter is required for 6-OHDA uptake into the CEP neurons, and inhibition of DAT-1 is protective from 6-OHDA induced degeneration (31, 32). *dat*-1 mutants were among the earliest to be identified as sensitized to swimming induced paralysis (SWIP), and synaptic localization of DAT-1 is required to prevent SWIP (59). Our observation of increased tendency towards SWIP after sugar exposure is consistent with modified dopamine transport, and is confirmed by the loss of the SWIP phenotype in sugar-fed *dat-1* KO worms. This could be an attempt to maintain dopamine in the synaptic cleft despite lower total dopamine levels, a possibility bolstered by recent evidence that high glucose diets decrease dopamine levels (55). Though little work has explored the relationship between DAT-1 and high sugar diets, we report a nearly 30% decrease in *dat-1* promoter-driven GFP fluorescence.

Though this decrease was smaller than that reported in recent work in *C. elegans* demonstrating an 80% decrease after high-glucose exposure, these combined findings support a relationship between high sugar diets and modulation of dopamine transmission in *C. elegans* (55). In mammalian models, high glucose diets activate protein kinase C, which drives DAT endocytosis, opening the possibility for a similar high sugar driven effect in worms (60–62). Notably, alterations to SWIP were not associated with increased neurodegeneration.

Beyond the toxicant-induced neurological impacts, we also deepen understanding of how the two most consumed sugars compare in their biological effects. High-glucose and high-fructose diets in our study produced similar but non-identical effects in nearly all experiments. Only glucose-fed worms exhibited elongated neuronal mitochondria, and in general, apart from lifespan, fructose-fed worms typically showed a less significant departure from controls than glucose-fed worms. These discrepancies may be explained by differences in the metabolism of these sugars, but it remains clear that the pathways driving alterations in dopaminergic function, lifespan, reproduction, and ATP production are impacted in very similar ways. This may indicate that in models with more complex organ systems, where stronger differences between sugar types have been observed, those differences are driven by tissue-specific metabolism and responses. For example, fructokinase, fructose bisphosphate aldolase-B, and dihydroxyacetone kinase, the three enzymes responsible for fructose metabolism, are only found in the liver and kidney of rats (63). In *C. elegans*, hexokinases (HXK-1,2,3) predicted to carry out the first step in fructose metabolism, are expressed ubiquitously (64). Thus, the reported mitochondrial swelling and respiratory dysfunction induced only by fructose in rat livers may be a result of specific mitochondrial dynamics and concentrated fructose metabolites in the liver and kidneys, versus the potentially non-tissue-specific metabolism in worms (65). It should be noted, in the same study in rats, high sugar diets had generally similar effects on fatty acid oxidation and mitochondrial protein acetylation in isolation, but divergent effects when supplemented on top of a high fat diet (65). Thus, further examination of dietary components in isolation and combination will be required to understand the complex dynamics governing effects of different sugars and how they relate to other model organisms.

The interaction between diet and toxicant exposure remains an active area of investigation due to the plethora of dietary alterations and chemicals that currently occur (66–71). High sugar diets elicit several metabolic and oxidative stress pathway alterations, depending on the exposure paradigm, leading to interactions with toxicants that target the same pathways. In *Drosophila melanogaster*, high glucose enhances Bisphenol A toxicity by exacerbating downregulation of testis-specific genes and upregulation of ribosome-associated genes (71). In *C. elegans*, high sugar diets increase susceptibility to monocrotophos and parathion, including increasing the damage inflicted on dopaminergic neurons (17, 18, 72). We show that both high-glucose and high-fructose decrease susceptibility to 6-OHDA induced neurodegeneration but enhance susceptibility to rotenone induced ATP depletion. Together, these data highlight the critical need to continue assessing toxicant by diet interactions for multiple endpoints, as the outcomes are likely highly specific to the tissue of interest and toxicokinetics of each chemical used.

There are also limitations to our study. We do not address how ecologically relevant neurotoxicants would interact with high sugar consumption. Due to the unique toxicokinetic effect of DAT-1 on 6-OHDA uptake, which would not generally be conserved for other dopaminergic toxicants, impacts of high sugar should be examined not only with ecologically relevant pollutants, but toxicants with varied mechanisms of toxicity. It is also possible the mechanisms we observe are limited to our exposure paradigm. As shown in Table 1, previous investigations often rely on developmental acute exposures and show different neurodegenerative results. Further work is required to understand the patterns of redox, bioenergetic, and dopaminergic transmission changes that occur both as a function of age and sugar consumption.

## Conclusions

Adult high-glucose and high-fructose diets are protective against 6-OHDA induced dopaminergic neurodegeneration, potentially due to their modifications of dopamine transmission processes decreasing 6-OHDA uptake. Intriguingly, this protection occurs despite decreased lifespan, decreased fecundity, and increased lipid storage. As demonstrated by the lack of neurodegeneration induced by ATP depletion, the induction of oxidative stress appears to be more important in the induction of dopaminergic neurodegeneration by 6-OHDA. This study highlights the important interactions between lifestyle factors such as diet, oxidative stress, and susceptibility to toxicant induced dopaminergic neurodegeneration.

## Materials and Methods

### Strains and Culture

The wild-type CGC (N2), LIU2 (Idrls[mdt-28p::mdt-28::mCherry + unc-76(+)]), BY200 (pdat-1:GFP), JMN080(pdat-1::MLS::GFP), SJ4103 (pmyo-3::mitoGFP), JV2 (*jrls2*[rpl-17p::Grx-1-roGFP2+unc-119(+)]), PE255 (*feIs5*[sur-5p::luciferase:GFP+rol-6(su1006)]), and PHX2923 (pdat-1::PercevalHR), PHX2867 (pdat- 1::MLS::roGFP), RM2702 (dat-1(ok157), were maintained at 20 C on K-agar plates seeded with OP50 *E. coli*. For experiments, worms were synchronized through egg-lays as in which worms were transferred onto new plates, allowed to lay eggs for 3 hours, then washed to remove adults. They were aged to adulthood on K-agar plates seeded with OP50 *E. coli*. As D1 adults (72 hours post egg lay), worms were evenly split between NGM, NGM + 100 mM glucose, or NGM + 100 mM fructose plates freshly seeded with OP50 E. coli. They were transferred daily to freshly seeded plates to discount effects of plate acidification. All worms were reared from D1-D5 of adulthood on their respective group plate (control, glucose, or fructose). On D8 assays were run, initiated, or worms were returned to K-agar OP50 plates as described for individual assays.

### Generation of transgenic strains

#### Generation of dat-1p::MLS::GFP

To generate the *dat-1*p::MLS::GFP plasmid, a 886 bp fragment directly upstream of the dat-1 start codon was amplified from wildtype (N2 Bristol type) genomic DNA, using primers with overhangs that contained homology to a plasmid containing a mitochondrial localization sequence (MLS) and GFP, Forward Primer 5’→3’: agggcgaattgggtaccCGTCTCATTCCTCATCTCCGAGC and Reverse Primer: 5’→3’: GTGCCATatcgatGGCTAAAAATTGTTGAGATTCGAGTAAACCG. The mitochondrial localization sequence was originally amplified from Fire Vector pPD96.32. pPD96.32 was a gift from Andrew Fire (Addgene plasmid # 1504 ; http://n2t.net/addgene:1504 ; RRID:Addgene_1504). The amplified dat-1 promoter was inserted into the plasmid containing the MLS and GFP using Gibson Assembly and insertion was confirmed by colony PCR with a nested GFP reverse primer and M13 Forward primer. The plasmid was sequenced to check for any mutations and co-injected with 50 ng/µl unc-119 rescue DNA, 50 ng/µl pBsSK, and 50 ng/µl EcoR1 cut salmon sperm DNA into unc-119(ed4) hermaphrodites. Once extrachromosomal lines were established and dat-1p::MLS::GFP signal was observed, plasmid was integrated by gamma irradiation as previously described (73). Integrated lines were outcrossed with N2 to remove possible background mutations.

#### Generation of dopaminergic neuron PercevalHR and mitochondrial roGFP

Worm strains expressing mitochondrial targeted reduction oxidation sensitive GFP (roGFP) and PercevalHR within the dopaminergic neurons were generated by SunyBiotech (https://www.sunybiotech.com). Both constructs were cloned into the pPD95.77 vector and included 890 bp of the *dat-1* promoter (ending just upstream of the start codon) amplified from genomic DNA and the 5’UTR from the *unc-54* gene present in the pPD95.77 vector. We inserted the reporter genes immediately downstream of the *dat-1* promoter and upstream of the *unc-54* 5’UTR. The mito-roGFP2- Grx1 coding sequence was adapted from pUAST mito roGFP2-Grx1 (Addgene Plasmid# 64995) by codon optimizing for expression in *C. elegans* using the *C. elegans* Codon Adapter (74) and a single intron was added 402 bp downstream of the start codon. The Perceval-HR coding sequence was adapted from pRsetB-his7-Perceval (Addgene Plasmid# 20336) by codon optimizing for *C. elegans* expression and a single intron was added 444 bp downstream from the start codon. The construction of the plasmids, verification by sequencing, microinjection into animals, integration, and isolation of individual strains were performed by SUNY Biotech. We received three low-copy-number strains for each construct, and all three strains were phenotypically normal.

### Fluorescence Microscopy

Strain and specific image analysis details are listed below for each individual endpoint. However, all strains were imaged with a Keyence BZX-2710 microscope. Unless otherwise specified, worms were transferred by stainless steel pick to 2% w/v agarose pads, anesthetized with 15-20 uL 0.5 M sodium azide, and imaged immediately. All quantitative image analysis was performed using Fiji ImageJ software. Background subtraction was performed for all quantitative microscopy by subtracting the equivalent measurement (mean grey value, etc) of a region in the image without a worm present.

### Fat Quantification

After 5 days of dietary exposure, LIU2 worms were washed off plates and washed three times with K- medium to remove bacterial debris. Worms were visualized in brightfield and under and EGFP or TexasRed filter with a 10X objective. Three independent experiments were conducted, with approximately 20 worms per treatment group imaged in each. Worm bodies were outlined as the region of interest (ROI) and mean grey value was determined within the full body of the worm.

### Lifespan

Lifespan assays were carried out with few modifications from previous descriptions (source). After 5 days of dietary exposure, 50 BY200 transgenic worms from each treatment group were transferred to 6 cm K-agar plates seeded with OP50. Worms were transferred to fresh plates every third to fourth day and monitored daily for death. Worms were considered dead if they displayed no touch response when poked with a steel pick and no bodily movement was observed. Animals that crawled off the plate or died due to vulval protrusion or bagging were censored. The data is represented as starting on day 8 as only worms that were alive on day 8 were used for subsequent analysis.

### Reproduction

Single worms were transferred to individual 6 cm NGM or NGM + sugar plates as late L4. They were transferred to a second plate 48 hours later, then were transferred every 24 hours to the remaining plates. Progeny were counted 48 hours after the adult was removed. At least 5 worms per treatment were utilized in each of three biological replicates.

### Dopaminergic Neurodegeneration

#### Exposure and Imaging

On day 5 of dietary exposure, worms were washed and dosed with 25 mM or 50 mM 6- hydroxydopamine (6-OHDA) in 10 mM or 20mM Ascorbic Acid (AA) solution [Ascorbic Acid, K+ mixture (K-medium, Cholesterol, CaCl2, MgSO4)]. Control groups were incubated in an identical volume of only the AA solution. All groups were incubated for 1 hour with rocking and subsequently washed with K- medium solution three times to ensure complete removal of 6-OHDA. They were replated on K-agar plates seeded with 2X OP50 and incubated at 20 C for 48 hours prior to imaging. Images were obtained with the 40X objective in Z-stacks encompassing the full head of the worm. Images were processed by generating maximal value Z-projections of stacks with FIJI ImageJ software and cropped to only display the head region of a single worm per image.

#### Scoring

Images were blindly scored with the open access software Blinder, by Solibyte solutions (75). Each dendrite of the four CEP neurons of each worm were scored on a scale of 0-4, as follows:

0−no visible damage or abnormalities
1−blebs or kinks encompassing less than 50% of the dendrite
2−blebs or kinks on encompassing more than 50% of the dendrite
3−breaks present with more than 50% of the dendrite remaining
4−breaks present with less than 50% of the dendrite remaining

To ensure scoring validity, the built-in Quality Control feature was utilized, and images were rescored until the error was less than 15%. Statistical significance was quantified by chi-squared test with a Bonferroni corrected p-value. Due to the high number of comparisons, letters are used to demonstrate results of pairwise comparisons. Statistically significant differences are represented by no overlapping letters when comparing two bars.

#### Seahorse Analysis

On day 5 of dietary exposure, whole worm respirometry was performed with a Seahorse Xf^e^24 Extracellular flux analyzer as previously described (76), with the following modifications: Worms were diluted to approximately 30 worms per well to maintain optimal oxygenation of the well during the protocol and 40 uM DCCD was utilized to obtain maximal inhibition of mitochondrial electron transport chain Complex V. 100-500 worms were reserved from each replicate and immediately imaged on K-agar plates for size determination. Worm volume was quantified using the WormSizer ImageJ plugin for normalization to worm volume.

### Mitochondrial Morphology

On Day 5 of adulthood, strains SJ4103 and JMN080 worms were imaged with the 60X and 40X objectives respectively. Images were taken as z-stacks encompassing the full muscle cell and maximally projected for analysis in ImageJ. CEP neuron mitochondria were analyzed for the number and length of each mitochondrion within each dendrite by using the line tool to manually trace each mitochondrion. Body wall muscle mitochondria were analyzed for mean grey value per muscle cell as a proxy for total mitochondrial mass.

### Swimming Induced Paralysis

On day 5 of the dietary exposure protocol approximately 10 worms were picked by flame sterilized steel pick into 100 uL of Millipore water in one well of a 96 well plate, each well was recorded for a minimum of 10 minutes from the time the pick was removed from the water. Videos were analyzed for the percentage of worms paralyzed at one-minute intervals for ten minutes, starting from the time the pick entered the water. Worms were considered paralyzed when they were completely rigid for 5 second intervals before and after the one-minute mark.

### Autofluorescence

On day 5 of dietary exposure, Bristol N2 worms were imaged with the 10X objective under brightfield, 405 nm, and 488 nm excitation. The brightfield image was used to select the entire body of the worm as the ROI and fluorescence was quantified in each respective channel as mean grey value.

### roGFP and PercevalHR imaging

On day 5 of dietary exposure, dat-1p::MLS::roGFP or JV2 worms, were paralyzed with 1 mM levamisole HCl and imaged. PercevalHR was mounted on 5% w/v agarose pads and imaged without paralytics. JV2 was imaged with the 10X objective and both dopaminergic strains were imaged with the 40X objective at 405 nm and 488 nm excitation. Mean grey value was quantified and compared in as a ratio or 405/488 for roGFPs and 488/405 for PercevalHR.

### qPCR

On day 5 of dietary exposure RNA was extracted via the Qiagen RNeasy Mini Kit (Qiagen 74104). Briefly, 100-200 worms were collected into conical tubes in K-medium. All animals were shaken for 10 minutes on an orbital shaker to clear gut bacteria, transferred to a 1.5 mL microcentrifuge tube, and suspended in RLT buffer. Samples were immediately flash frozen in liquid nitrogen and thawed on ice. Disruption of the worm cuticle was then completed by bead beating with zirconia beads for 8 cycles of 30 seconds beating and 1 minute on ice. Homogenate was then utilized in accordance with kit instructions. cDNA was synthesized from 2 ug total RNA with a high-capacity cDNA Reverse Transcription Kit in 20 uL reactions (Thermo Fischer, Ref. 4368814). We carried out qPCR using diluted cDNA, Power SYBR Greeen Master Mix (Thermo Fischer 4368702) and 0.5 μM of gene-specific primers (Sup Table 1) in a CFX96 qPCR Real-Time PCR module with C1000 Touch Thermal Cycler (BioRad). After 10 minutes at 95°C, a two-step cycling protocol was used (15 seconds at 95°C, 60 seconds at 60°C) for 40 cycles. We calculated relative expression using the ΔΔCt method with *cdc-42* and *tba-1* as reference genes. Genes only expressed in the dopaminergic neurons, *dat-1*, *dop-3*, and *cat-2*, required an additional 10 cycles of PCR amplification prior to qPCR for adequate detection. Three samples were recorded for each treatment for each biological replicate. Three biological replicates were performed. Any data point for which the standard deviation between three technical replicates was not below 0.300 was discarded. Due to the low transcript levels of some antioxidant genes, fewer samples were valid for these genes.

### Total GSH levels

On day 5 of dietary exposure, 200 worms from each group were suspended in 75 uL of MES buffer in 1.7 mL conical tubes. Approximately 50 uL 0.5 mm zirconium oxide beads were added to each tube. Samples were homogenized via bead beating (8 cycles, 30s on sonication, 30s without sonication at 4 C). Next, 65 uL of homogenate was recovered, with 10 uL transferred to a new tube for protein quantification and 55 uL diluted by half with 5% w/v metaphosphoric acid for total GSH quantification. GSH quantification was completed in accordance with instructions for tissue homogenate (Cayman Chemical Kit No.703002). Protein quantification for each sample was conducted by BCA Assay in accordance with kit instructions (Millipore Sigma 71285M).

### Whole Worm ATP Levels

Strain PE255 worms were reared in accordance with the dietary exposure protocol, challenged with 20 uM rotenone for 1 hour, and transferred to 96-well plates for Luciferin-based ATP quantification as previously described (77). Luminescence was normalized to number of worms per well rather than GFP due to autofluorescence differences.

### p*dat-1*::GFP fluorescence quantification

Strain BY200 worms were reared in accordance with the dietary exposure protocol. On day 8, worms were picked onto 2% w/v agarose slides and paralyzed with 60 mM sodium azide. Images of the head region of each worm were taken in z-stacks with 0.5 uM pitch such that the full cell body was captured. Z-stacks were maximum projected and assessed for mean grey value in FIJI ImageJ software.

### Graphing and Statistical Analysis

GraphPad Prism version 9.5.0 was used for all graph generation and statistical testing. Statistical tests are identified in the figure legend of each graph.

Supplementary Figure 1. Survival of worms during the 5-day exposure to high sugar. BY200 worms were reared to Day 1, at which point 20 worms per treatment per replicate were transferred to control NGM plates or plates containing 100 mM glucose or 100 mM fructose. Each day worms were assessed for survival by touch response with a flame sterilized platinum pick. Surviving worms were transferred to freshly seeded plates each day to avoid plate acidification and to separate them from progeny. For each group 3 biological replicates were assessed, with 20 individuals per replicate. A two-way ANOVA was performed to assess statistical significance, with no differences identified.

Supplementary Figure 2. P-values for individual chi-squared tests for analysis of dopaminergic neurodegeneration. Chi-squared tests were used to determine the outcome of 15 comparisons with a Bonferroni corrected p-value of 0.0033. Within each diet, the vehicle was compared to both doses of 6- OHDA. Across diets each treatment was compared. Results are color coded for clarity: grey was not assessed, green is statistically significant, yellow is not statistically significant

Supplementary Figure 3. Muscle cell mitochondrial mean grey value. SJ4103 worms were reared in accordance with the dietary exposure protocol and imaged on day 8. Images were obtained using a Keyence BZ-X710 with 60X magnification (oil immersion). Z-stacks were set to encompass the entirety of the cell, and maximum projected for analysis. Individual cells were outlined as the region of interest for analysis in Image J. Mean grey value was used as a proxy for total mitochondrial area (One-way ANOVA).

Supplementary Figure 4. Oxygen consumption rate normalized to worm number without accounting for worm size. Whole worm respirometry was performed with the Seahorse XF24 Bioanalyzer and reported as oxygen consumption rate normalized to the number of worms in each well. All wells from the same biological replicate were averaged to produce n=1 (One-way ANOVA, Tukey’s Post Hoc, *p<0.0332).

Supplementary Figure 5. Autofluorescence quantification at 488 nm and 405 nm excitation wavelengths. N2 (wild-type) worms were reared on their respective sugar diets and imaged at day 5 of adulthood to determine if high sugar diets alter worm autofluorescence. No significant difference was detected in the N2 strain at 488 nm, however both high glucose and high fructose diets increase autofluorescence at 405 nm excitation (One way ANOVA, Tukey’s Post Hoc,**p<0.0021,****p<0.0001)

Supplementary Figure 6. Relative gene expression of individual genes.

Supplementary Figure 7. Swimming Induced Paralysis Timecourse. BY200 worms were synchronized by timed egg lay and assessed for susceptibility to swimming induced paralysis at times correlating to various developmental and reproductive stages: 24-hours (L2 larval stage), 48-hours (L4 larval stage), 72-hours (early adults), 120-hours (mid-reproductive age adults), 168-hours (late-reproductive age adults), and 192-hours (post-reproductive, experiment timepoint). Three biological replicates were assessed for 24-168 hours, with 2 wells containing approximately 10 individuals each per replicate (n=6 per strain, per timepoint). The data for 192-hours is the data represented for D8 in figure 6D. Results were assessed by two-way ANOVA followed by Sidak’s Test for multiple comparisons with (p<0.05) as the threshold for significance. Only the 48-hour (L4) timepoint indicated a significant difference, p<0.0001.

**Supplementary Table 1.**
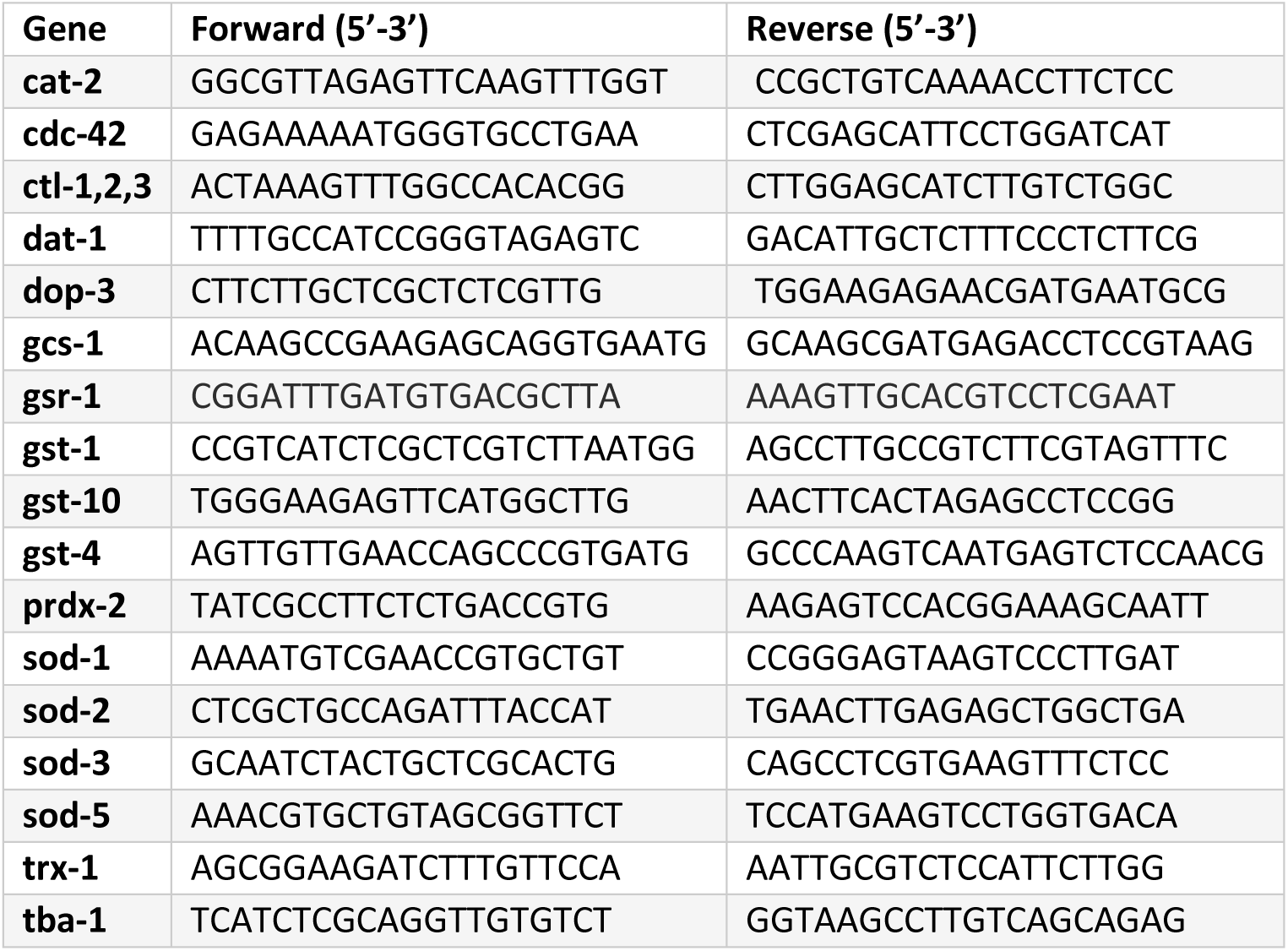
Primers utilized for RT-qPCR.

## Availability of Data and Materials

*C. elegans* strains JMN080, PHX2923, and PHX2867 are available from the corresponding author upon request. All other strains are available from the *C. elegans* Genome Center. All data and the source video and images are available upon request.

## Acknowledgements

This work was funded by NIH awards K99-ES029552 (JHH), NIEHS P42ES010356 (JNM), NIEHS T32ES021432 (KSM), and R01ES034270 (JNM), and strains were provided by the Caenorhabditis Genetics Consortium.

