## Supplementary figures and images for "Chronic high-sugar diet in adulthood protects *Caenorhabditis elegans* from 6-OHDA induced dopaminergic neurodegeneration"

### Supplemental Figure 1

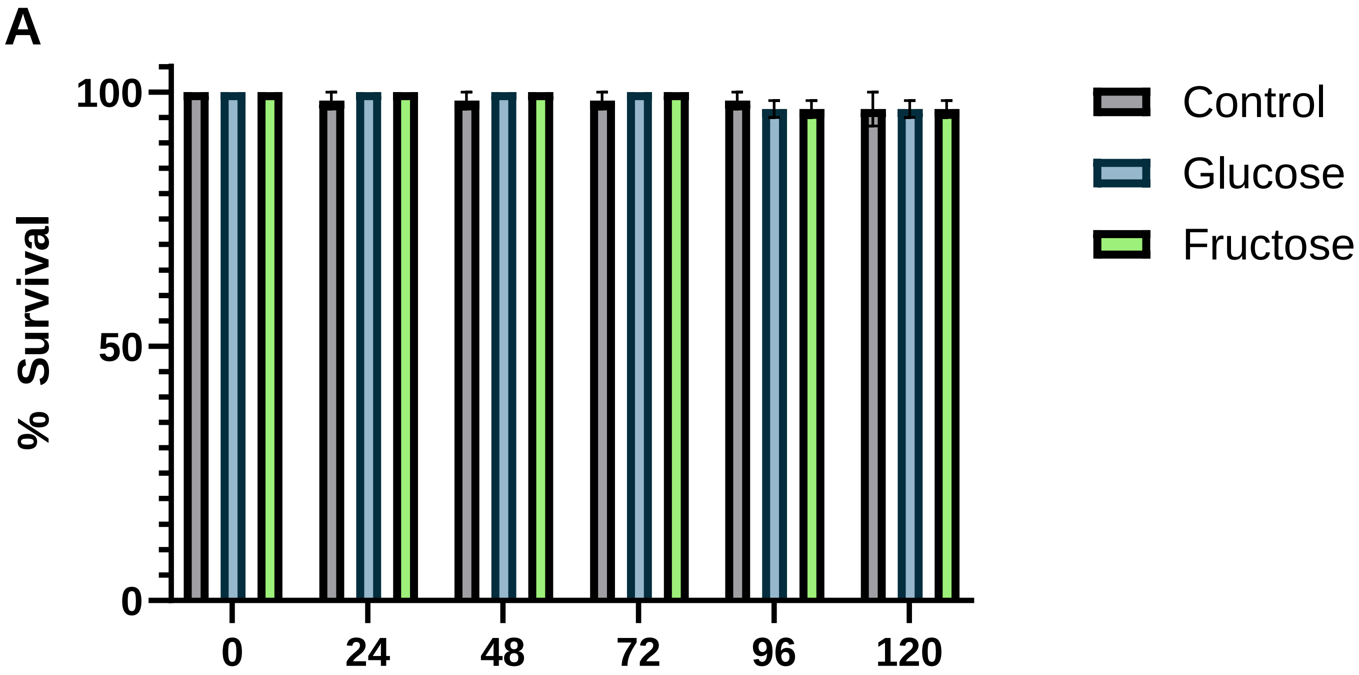

### Supplemental Figure 3

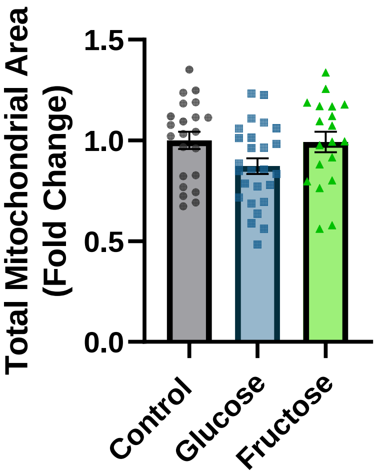

### Supplemental Figure 4

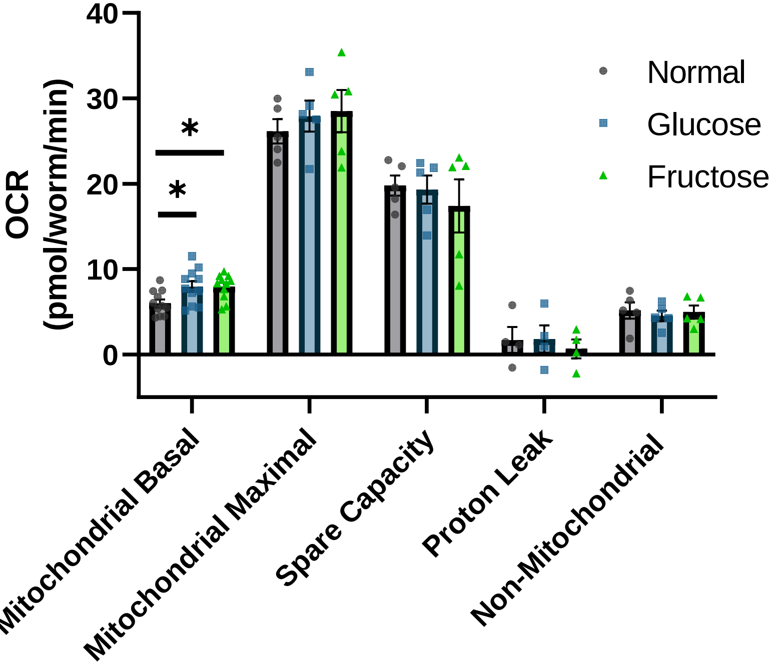

### Supplemental Figure 5

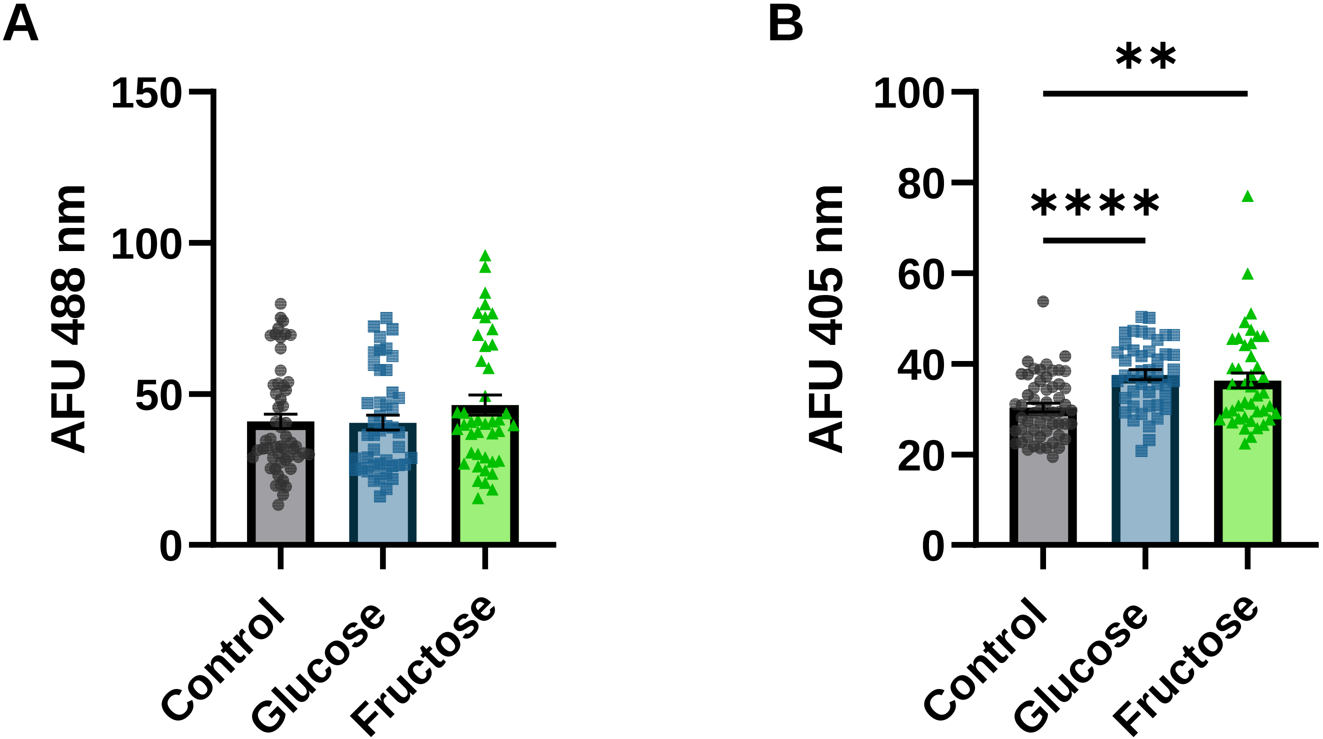

### Supplemental Figure 6

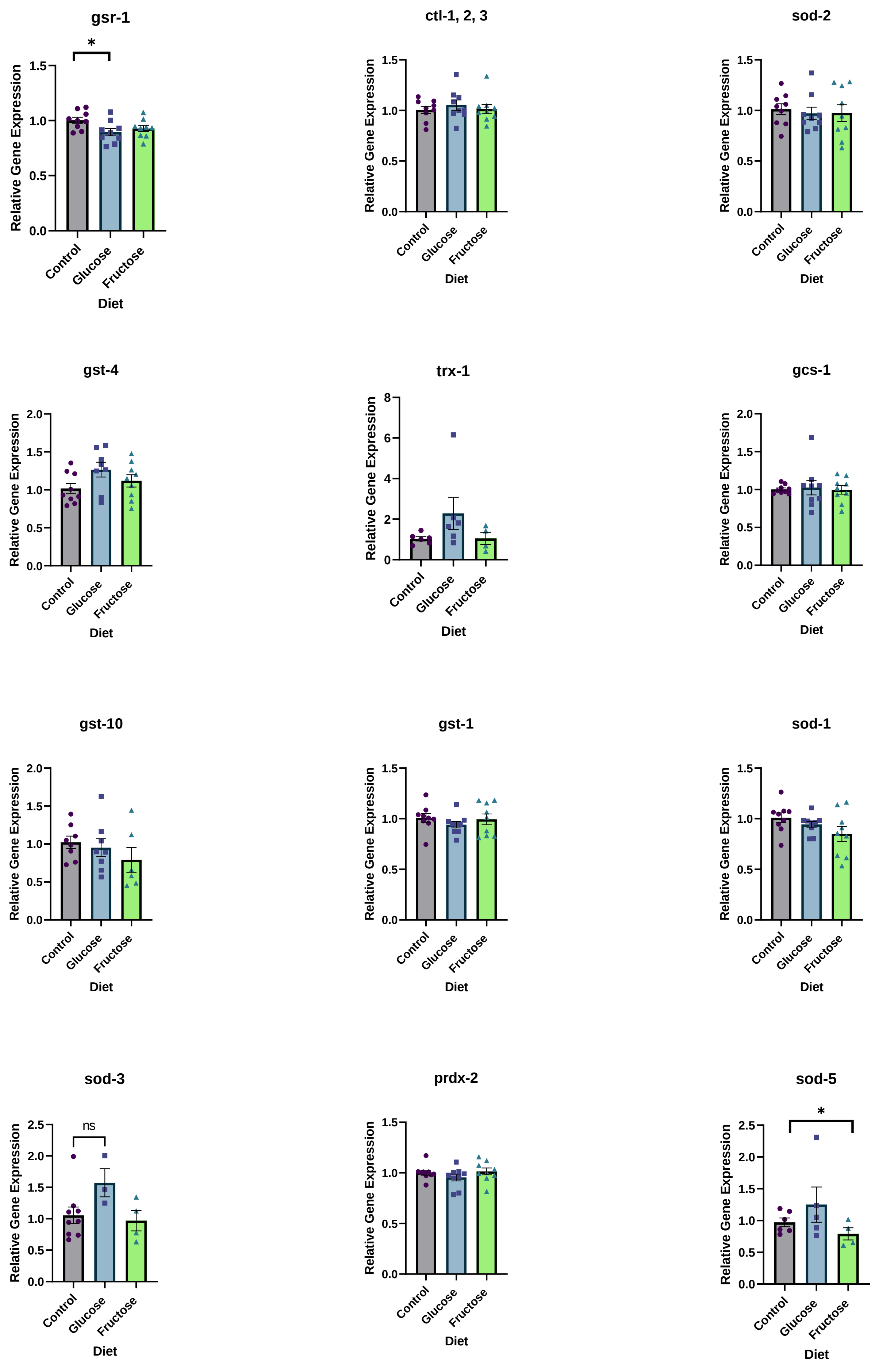

### Supplemental Figure 7

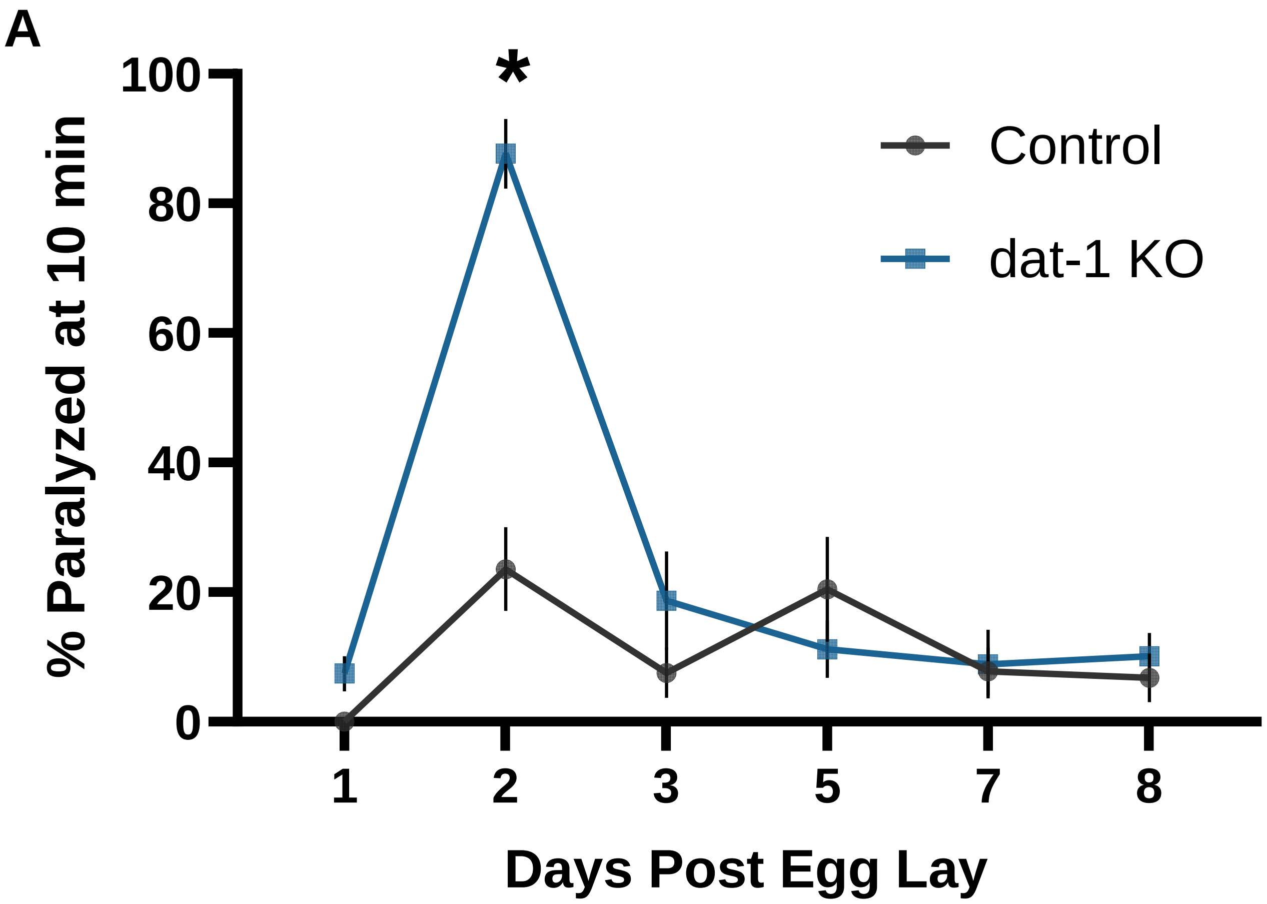
