## Supplemental Figure 2 for "Chronic high-sugar diet in adulthood protects *Caenorhabditis elegans* from 6-OHDA induced dopaminergic neurodegeneration"

| Bonferroni corrected p-value: 0.00333 | | Control | | | Glucose | | | Fructose | | |
|---|---|---|---|---|---|---|---|---|---|---|
| | | Vehicle | 25 mM 6-OHDA | 50 mM 6-OHDA | Vehicle | 25 mM 6-OHDA | 50 mM 6-OHDA | Vehicle | 25 mM 6-OHDA | 50 mM 6-OHDA |
| Control | Vehicle | | 0.0010 | <0.0001 | 0.4706 | | | 0.4706 | | |
| | 25 mM 6-OHDA | | | | | 0.0027 | | | 0.0195 | |
| | 50 mM 6-OHDA | | | | | | <0.0001 | | | <0.0001 |
| Glucose | Vehicle | | | | | 0.0021 | <0.0001 | 0.4706 | | |
| | 25 mM 6-OHDA | | | | | | | | 0.3798 | |
| | 50 mM 6-OHDA | | | | | | | | | 0.1120 |
| Fructose | Vehicle | | | | | | | | 0.0229 | <0.0001 |
| | 25 mM 6-OHDA | | | | | | | | | |
| | 50 mM 6-OHDA | | | | | | | | | |
